# Exploiting the *ZIP4* homologue within the wheat *Ph1* locus has identified two lines exhibiting homoeologous crossover in wheat-wild relative hybrids

**DOI:** 10.1101/142596

**Authors:** María-Dolores Rey, Azahara C Martín, Janet Higgins, David Swarbreck, Cristobal Uauy, Peter Shaw, Graham Moore

## Abstract

Despite possessing related ancestral genomes, hexaploid wheat behaves as a diploid during meiosis. The wheat *Ph1* locus promotes accurate synapsis and crossover of homologous chromosomes. Interspecific hybrids between wheat and wild relatives are exploited by breeders to introgress important traits from wild relatives into wheat, although in hybrids between hexaploid wheat and wild relatives, which possess only homoeologues, crossovers do not take place during meiosis at metaphase I. However, in hybrids between *Ph1* deletion mutants and wild relatives, crossovers do take place. A single *Ph1* deletion (*ph1b*) mutant has been exploited for the last 40 years for this activity. We show here that selection of chemical induced mutant lines possessing mutations in *TaZIP4-B2* exhibit high levels of homoeologous crossovers when crossed with a wild relative. Exploitation of *Tazip4-B2* mutants rather than mutants with whole *Ph1* locus deletions may improve introgression of wild relative chromosome segments into wheat. Such mutant lines may be more stable over multiple generations, as multivalents causing accumulation of chromosome translocations are less frequent.

**Key message:** Exploiting the *ZIP4* homologue within the wheat *Ph1* locus has identified two wheat mutants through a non-GM route, which can be exploited as an alternative to the Chinese Spring *ph1b* mutant in wheat introgression strategies.

## Introduction

Hexaploid bread wheat (*Triticum aestivum*) is composed of three related ancestral genomes (A, B and D), each containing 7 identical (homologous) chromosome pairs (homologue pairs 1-7). Each homologue pair has a corresponding related homoeologous chromosome pair, possessing similar gene order and content, within each of the other two genomes. Despite the similarity between homoeologues, wheat behaves as a diploid at meiosis, with synapsis and crossovers (COs) only occurring between homologous chromosomes, rather than between homoeologous chromosomes (for example 1A only pairs with 1A but not with either 1B or 1D, holding true for all seven chromosome groups). This diploid-like behaviour is predominantly controlled by *Ph1*, a dominant locus on chromosome 5B (Riley and Chapman 1958; Sears and Okamoto 1958). Sexual hybridisation between wheat and wild relatives (for example rye or *Aegilops variabilis*) produces interspecific hybrids, containing haploid sets of wheat and wild relative homoeologous chromosomes, but exhibiting virtually no COs during meiosis. However, in *Ph1* deleted wheat-rye hybrids, an average of 7 COs per cell is observed (Sears 1977; Dhaliwal et al. 1977). Such interspecific hybrids have been used to introgress important traits from wild relatives into wheat.

Synapsis is a process early in meiosis by which homologues intimately align with each other, forming bivalents held together by the proteinaceous structure or synaptonemal complex (SC). Ultimately the SC is degraded, so that the bivalents are then held together by chiasmata or COs at metaphase I, allowing their correct segregation. Recently, *Ph1* has been shown to have a dual effect on synapsis and CO formation in wheat (Martín et al. 2014; Martín et al. 2017). The effect on synapsis occurs during the telomere bouquet stage, when *Ph1* promotes more efficient homologous synapsis, thereby reducing the chance of homoeologous synapsis (Martín et al. 2017). The effect on CO formation happens later in meiosis, when *Ph1* prevents MLH1 sites (Double Holliday Junctions marked to become COs) on synapsed homoeologues from becoming COs. The *Ph1* locus was defined as a deletion phenotype, first described by scoring meiosis in wheat hybrids lacking the whole 5B chromosome (Riley and Chapman 1958; Sears and Okamoto 1958). Exploitation of smaller chromosome 5B deletions, later characterised and defined the *Ph1* locus to a region on chromosome 5B, containing a duplicated chromosome 3B segment carrying heterochromatin and *TaZIP4-B2* (originally termed *Hyp3*, UniProtKB - Q2L3T5), inserted into a cluster of *CDK2*-like genes interspersed with methyl transferase genes (originally termed *SpG*, UniProtKB-Q2L3W3) (Griffiths et al. 2006; Al-Kaff et al. 2008; Martín et al. 2017). Ethylmethane sulphonate (EMS) treatment failed to produce mutants exhibiting the full *Ph1* deletion phenotype (Griffiths et al. 2006). This observation, combined with the dual activity of the *Ph1* locus on synapsis and COs, suggested that the *Ph1* phenotype was likely to result from the activity of more than a single gene (Griffiths et al. 2006).

A single *Ph1* deletion mutant, developed in hexaploid wheat variety Chinese Spring (CS) (CS *ph1b*), has been used by breeding programmes worldwide to introgress wild relative chromosome segments into wheat. However, the CS *ph1b* mutant is reported to accumulate extensive rearrangements reducing fertility (Sánchez-Morán et al. 2001), due to homoeologous synapsis and COs as visualised by the occurrence of multivalents at metaphase I during meiosis. It would therefore be useful to identify novel wheat *Ph1* mutant lines, with reduced homoeologous synapsis and CO at meiosis, but which do exhibit homoeologous COs in hybrids with wild-relatives. As described previously, the *Ph1* locus, which affects both synapsis and CO, is a complex cluster of *CDK2*-like and methyl transferase genes containing a *ZIP4* paralogue. It has been proposed that *Ph1*’s effect on synapsis is connected to altered Histone H1 CDK2-dependent phosphorylation in the presence and absence of *Ph1*. Altered phosphorylation affects chromatin structure and delays premeiotic replication, subsequently affecting homologue synapsis, thus allowing homoeologous synapsis to take place (Greer et al. 2012; Martín et al. 2017). Lines carrying mutations in the *Ph1 CDK2*-like homologue in Arabidopsis, also exhibit reduced synapsis under specific conditions, suggesting a role for these genes in efficient synapsis (Zheng et al. 2014). We have also previously proposed that the effect of *CDK2*-like genes on chromatin structure not only affects synapsis, but might also affect the resolution of Double Holliday Junctions (marked by MLH1) as COs (Greer et al. 2012). Okadaic acid treatment affects chromatin structure and can induce homoeologous CO in wheat-wild relative hybrids (Knight et al. 2008). However, given the locus contains multiple copies of the *CDK2*-like and methyltransferase genes, it would be a complex and laborious study to identify EMS mutants within these genes and combine possible mutations. The transfer of multiple mutated genes into different elite genetic backgrounds by breeders, for subsequent crossing with wild relatives, would also be laborious. Moreover, as previously indicated, we want to identify wheat mutant lines which exhibit reduced homoeologous synapsis or multivalents at metaphase I, so the *CDK2*-like genes would not be the initial candidates for such an approach. Although there are *ZIP4* homologues on group 3 chromosomes, the *ZIP4* paralogue (*TaZIP4-B2*) within the *Ph1* locus on chromosome 5B is single copy, compared to the *CDK2*-like and methyl transferase gene cluster. Moreover, *ZIP4* has been shown to have a major effect on homologous COs, but not on synapsis, in both Arabidopsis and rice (Chelysheva et al. 2007; Shen et al. 2012). Knockouts of this gene in diploids usually result in sterility, as elimination of homologous COs leads to metaphase I pairing failure and incorrect segregation. We therefore assessed whether the selection and scoring of *Tazip4-B2* EMS mutants would identify wheat lines with minimal homoeologous synapsis and CO (as observed by the occurrence of multivalents at metaphase I), but which exhibit homoeologous COs in hybrids with wild-relatives. We describe the identification of two such lines through this approach.

## Materials and methods

### Plant material

Plant material used in this study included: wild-type hexaploid wheat (*Triticum aestivum* cv. Chinese Spring and cv. Cadenza); a Chinese Spring mutant lacking the *Ph1* locus (*ph1b*); two *Tazip4-B2* mutant Cadenza lines (Cadenza1691 and Cadenza0348); hexaploid wheat- *Aegilops variabilis* hybrids - crosses between hexaploid wheat (*T. aestivum* cv. Cadenza) and *Ae. variabilis* (2n=4x=28) - using either wild type Cadenza or *Tazip4*-B2 mutant lines.

For meiotic studies, the seedlings were vernalised for 3 weeks at 8°C and then transferred to a controlled environmental room until meiosis under the following growth conditions: 16 h: 8 h, light: dark photoperiod at 20°C day and 15°C night, with 70% humidity. Tillers were harvested after 6 to 7 weeks, when the flag leaf was starting to emerge, and anthers collected. For each dissected floret, one of the three synchronised anthers was squashed in 45% acetic acid in water and assigned to each meiotic stage by observation under a LEICA DM2000 microscope (LeicaMicrosystems, <u>http://www.leica-microsystems.com/</u>). The two remaining anthers were either fixed in 100% ethanol: acetic acid 3:1 (v/v) for cytological analysis of meiocytes or frozen in liquid nitrogen and stored at −80°C for RNA-seq analysis.

### RNA-seq experiments

#### Sample preparation

Anthers from wild-type wheat (WT) and wheat lacking the *Ph1* locus (*ph1b* deletion) were harvested as described in the plant material section. Anthers at late leptotene-early zygotene stage were later harvested into RNA later (Ambion, Austin, TX). The anthers from three plants of each genotype were pooled in a 1.5 ml Eppendorf until reaching 200 to 400 anthers. Once sufficient anthers had been collected, the material was disrupted using a pestle, centrifuged to eliminate the RNA, and then homogenized using QIAshredder spin columns (Qiagen, Hilden, Germany). RNA extraction was performed using a miRNeasy Micro Kit (Qiagen, Hilden, Germany) according to the manufacturer’s instructions. This protocol allows purification of a separate miRNA-enriched fraction (used for further analysis) and the total RNA fraction (□200 nt) used in this study. This process was repeated to obtain three biological samples of each genotype.

#### RNA-seq library preparation and sequencing

1 μg of RNA was purified to extract mRNA with a poly-A pull down using biotin beads. A total of six libraries were constructed using the NEXTflex™ Rapid Directional RNA-Seq Kit (Bioo Scientific Corporation, Austin, Texas, USA) with the NEXTflex™ DNA Barcodes – 48 (Bioo Scientific Corporation, Austin, Texas, USA) diluted to 6 μM. The library preparation involved an initial QC of the RNA using Qubit DNA (Life technologies, CA, Carlsbad) and RNA (Life technologies, CA, Carlsbad) assays as well as a quality check using the PerkinElmer GX with the RNA assay (PerkinElmer Life and Analytical Sciences, Inc., Waltham, MA, USA). The constructed stranded RNA libraries were normalised and equimolar pooled into one final pool of 5.5 nM using elution buffer (Qiagen, Hilden, Germany, Hilden, Germany). The library pool was diluted to 2 nM with NaOH, and 5 μl were transferred into 995 μl HT1 (Illumina) to give a final concentration of 10 pM. 120 μl of the diluted library pool were then transferred into a 200 μl strip tube, spiked with 1% PhiX Control v3, and placed on ice before loading onto the Illumina cBot. The flow cell was clustered using HiSeq PE Cluster Kit v4, utilising the Illumina PE_HiSeq_Cluster_Kit_V4_cBot_recipe_V9.0 method on the Illumina cBot. Following the clustering procedure, the flow cell was loaded onto the Illumina HiSeq2500 instrument following the manufacturer’s instructions. The sequencing chemistry used was HiSeq SBS Kit v4 with HiSeq Control Software 2.2.58 and RTA 1.18.64. The library pool was run in a single lane for 125 cycles of each paired end read. Reads in bcl format were demultiplexed based on the 6 bp Illumina index by CASAVA 1.8, allowing for a one base-pair mismatch per library, and converted to FASTQ format by bcl2fastq.

#### RNA-seq Data Processing

The raw reads were processed using SortMeRNA v2.0 (Kopylova et al. 2012) to remove rRNA reads. The non rRNA reads were then trimmed using Trim Galore v0.4.1 (http://www.bioinformatics.babraham.ac.uk/projects/trim_galore/) to remove adaptor sequences and low quality reads (-q 20 --length 80 --stringency 3). 273,739 transcripts (Triticum_aestivum_CS42_TGACv1_scaffold.annotation) were quantified using kallisto v0.43.0 (Bray et al. 2016). The index was built using a k-mer length of 31, then Kallisto quant was run using the following options -b 100 --rf-stranded. Transcript abundance was obtained as Transcripts Per Million (TPM) for each gene.

### Cytological analysis and image processing

A total of five plants per line were examined. Anthers from *Tazip4-B2* Cadenza mutant lines, wild type Cadenza, *Tazip4-B2* mutant line*-Ae variabilis* hybrids and Cadenza-*Ae. variabilis* hybrids were harvested as described in the plant material section. Cytological analysis of meiocytes was performed using Feulgen reagent as previously described (Sharma and Sharma 1980). Images were collected using a LEICA DM2000 microscope (LeicaMicrosystems, http://www.leica-microsystems.com/), equipped with a Leica DFC450 camera and controlled by LAS v4.4 system software (Leica Biosystems, Wetzlar, Germany). Images were processed using Adobe Photoshop CS5 (Adobe Systems Incorporated, US) extended version 12.0 × 64.

### Nucleotide analysis

The regions of each mutation in both mutant Cadenza lines were sequenced to confirm the existence of either missense or nonsense mutations in Cadenza1961 and Cadeza0348 respectively. Wheat leaf tissues from wild type Cadenza and mutant Cadenza lines were harvested at growth stages 3-4 (Feekes scale). DNA was extracted using the CTAB method (Murray and Thomson 1980). Two pairs of primers were designed using the Primer3plus software (Untergasser et al. 2007) based on the *ZIP4* sequence called TRIAE_CS42_5BL_TGACv1_404600_AA1305800 (http://plants.ensembl.org/Triticum_aestivum/Info/Index). The primers used were: forward primer: 5´GCCGCCATGACGATCTCCGAG3´ and reverse primer: 5´GGACGCGAGGGACGCGAG3´ for Cadenza1691 and forward primer: 5´GTGTTCCTAATGCTCACAACTC3´ and reserve primer: 5´ACCAGACATACTTGTGCTTGGT3´ for Cadenza0348. PCR amplification was performed using MyFi Polymerase (Bioline Tauton, MA, USA), according to the manufacturer’s instructions. The primers were amplified as follows: 3 min 95 °C, 35 cycles of 15s at 95 °C, 15s at 58 °C and 30s at 72 °C. PCR products were resolved on 2% agarose gels in 1xTBE and stained with ethidium bromide and visualised under UV light. PCR products were purified using Qiaquick PCR Purification kits (Qiagen, Hilden, Germany) and cloned using a p-GEM T easy vector kit (Promega, Madison, Wisconsin, USA). The ligation mixture was used to transform *Escherichia coli* DH5a, and transformants were selected on LB agar containing ampicillin (100mg/ml) (Sigma, St. Louis, MO, USA), Isopropyl β-D-1-thiogalactopyranoside (IPTG, 100 mM) (Sigma, St. Louis, MO, USA), and 5-bromo-4-chloro-3-indolyl β-D-galactopyranoside (X-Gal, 20 mg/ml) (Sigma, St. Louis, MO, USA). The PCR fragments were isolated using QIAprep Spin Miniprep kit (Qiagen, Hilden, Germany). All kits were used as described in the manufacturer’s instructions. Sequencing was carried out by the Eurofins Company. Alignment of sequences was carried out using Clustal Omega software (<u>http://www.ebi.ac.uk/Tools/msa/clustalo/</u>).

### Statistical analyses

Statistical analyses were performed using STATISTIX 10.0 software (Analytical Software, Tallahassee, FL, USA). *Tazip4-B2* mutant line-*Ae. variabilis* hybrids were analysed by the Kruskal–Wallis test (nonparametric ANOVA) to compare the chiasma frequency among lines followed by Dunn’s Test, P < 0.05. *Tazip4-B2* mutant Cadenza lines were analysed by analysis of variance (ANOVA) based on randomised blocks. Several data transformations were applied to meet the requirement of homogeneity of variances. These included: exponential (univalents), cosine (rod bivalents and chiasma frequency) and sine (ring bivalents) transformations. Means were separated using the Least Significant Difference (LSD) test and with a probability level of 0.05.

## Results and Discussion

### *TaZIP4-B2* expression

In hexaploid wheat, *ZIP4* homologues are located within the *Ph1* locus on 5B, and also on chromosomes 3A, 3B and 3D. Before undertaking the targeted induced lesion in genomes (tilling) mutant analysis, we assessed the expression of *TaZIP4-B2* to confirm: that the *TaZIP4-B2* gene within the *Ph1* locus was expressed during meiosis; that it had a higher level of expression than those present on chromosome group 3; and finally, that deletion of *Ph1* significantly reduced overall *ZIP4* expression. At the coding DNA sequence and amino acid level, *TaZIP4-B2* (AA1305800.1) showed 95.3% and 89.2% similarity to *TaZIP4-B1* (AA0809860.1), 94.1% and 87.5% to *TaZIP4-A1* (AA0645950.1), and 94.4% and 87.2% to *TaZIP4-D1* (AA0884100.1) respectively (Supplementary Fig. 1). To compare the relative expression of these *ZIP4* homologues on chromosomes 3A, 3B, 3D and 5B, RNA samples were collected from anthers of hexaploid wheat (*Triticum aestivum* cv. Chinese Spring) (WT) and the *Ph1* deletion mutant (*ph1b*) at late leptotene-early zygotene stage, and six libraries were prepared for the RNA-seq study. RNA-seq analysis showed that *TaZIP4-B2* exhibited a higher level of expression than the *ZIP4* homologues on homoeologous group 3 chromosomes (Fig. 1; Supplementary Fig. 1). Moreover, *TaZIP4-B2* also showed three splice variants (Supplementary Fig. 2) in contrast to homoeologous group 3 chromosome *ZIP4* homologues. One of these splice variants (splice variant 1) accounted for 97% of the *TaZIP4-B2* transcripts. As expected, when the *Ph1* locus was deleted, the level of expression of *TaZIP4-B2* was also eliminated (*p < 0.05*), but there was no apparent increase in the transcription of the *ZIP4* homologues on homoeologous group 3 chromosomes, to compensate for the absence of *ZIP4* on chromosome 5B (*p > 0.05*) (Fig. 1; Supplementary Table 1). Thus, RNA-seq data revealed that the expression of *ZIP4* was derived mainly from the gene (*TaZIP4-B2*) on chromosome 5B, within the *Ph1* locus.

**Fig. 1.**
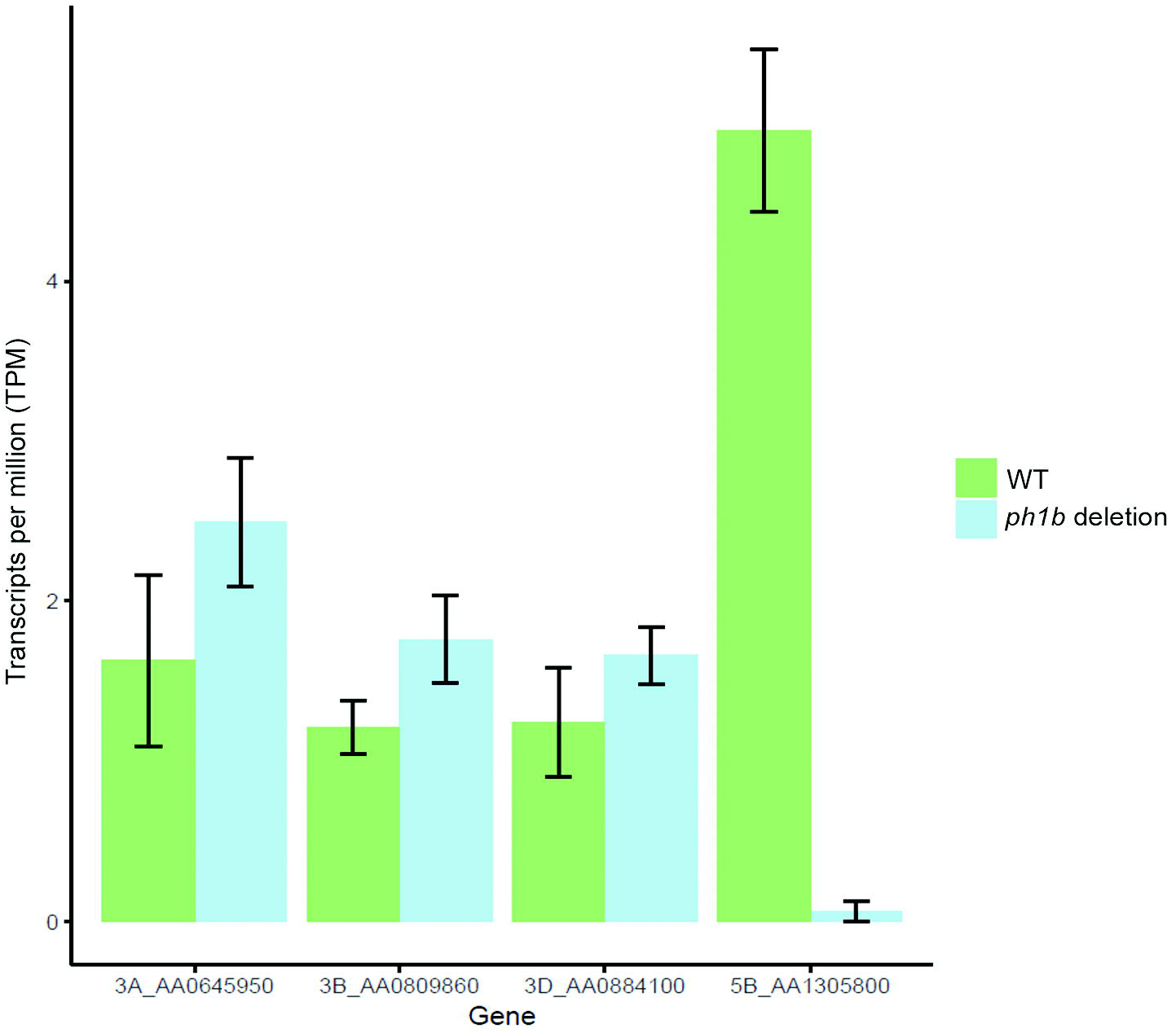
Relative expression of *ZIP4* homologs in *Triticum aestivum* cv. Chinese Spring in presence (WT) and in absence (*ph1b* deletion) of the *Ph1* locus obtained by RNA-seq analysis.

### *TaZIP4-B2* suppresses homoeologous COs in wheat-*Ae. variabilis* hybrids

The protein-coding sequences of 1,200 EMS mutant lines (Rakszegi et al. 2010) from hexaploid variety “Cadenza” have been recently sequenced using exome-capture, and displayed to allow the identification of millions of mutations in the sequenced genes (<u>www.wheat-tilling.com</u>) (Krasileva et al. 2017). The mutations identified are accessible using the wheat survey sequence (Mayer et al. 2014) via a database, which includes their location within the gene and the predicted effect that each variant has on its protein. Simply searching this database reveals those plants possessing mutations in the target genes, as well as a list of all mutations possessed by the plant (Krasileva et al. 2017). We selected seven of the 1,200 EMS mutant lines, which possessed potentially interesting mutations within *TaZIP4-B2* (Traes_5BL_9663AB85C.1) (Supplementary Table 2). Five of these mutant lines exhibited regular pairing at meiotic metaphase I, so were not taken further. However, two of the mutant lines (Cadenza1691 and Cadenza0348) showed reduced number of COs, suggesting that their *Tazip4-B2* mutations exhibit a phenotype. Both lines were selected for wide crossing studies with wild relatives to score the effect of their *Tazip4-B2* mutations on homoeologous CO frequency in the wheat *Tazip4-B2* mutant-wild relative hybrids, as compared to wheat wild type-wild relative hybrids. Mutations within *TaZIP4-B2* were verified by sequencing, and primers were designed to the mutated regions to follow mutated genes during crossing (Supplementary Fig. 3 and Materials and Methods). *Tazip4-B2*, within the Cadenza1691 mutant line, possessed a missense mutation within one of the tetratricopeptide repeats (C to T change leading to an A167V), shown to be important for ZIP4 function (Perry et al. 2005). *Tazip4-B2*, within the Cadenza0348 mutant line, possessed a nonsense mutation (a premature stop codon: G to A change leading to W612*) (Supplementary Fig. 3). In addition to these *Tazip4-B2* mutations, the two mutant lines also possessed mutations (mostly missense, but also splice or stop codons) within the coding sequences of 106 other shared genes. Sixteen of these genes, including *TaZIP4-B2*, were located on chromosome 5B. However, none of the genes apart from *TaZIP4-B2* were located within the 2.5 MB *Ph1* region defined in our previous study (Griffiths et al. 2006; Al-Kaff et al. 2008).

Compared to the chromosome 5B deletion mutant - wild relative hybrid, no other wheat chromosome deletion mutants have previously been reported as exhibiting a similar level of homoeologous CO formation at metaphase I (Riley and Chapman 1958; Sears 1977). For example, the 3D locus *Ph2*, exhibits a four-fold lower level of induction compared to *Ph1* (Prieto et al. 2005). Equally, deleting regions of chromosome 5B apart from the 2.5 MB *Ph1* region did not result in homoeologous CO formation at metaphase I when the lines were crossed with wild relatives (Roberts et al. 1999; Griffiths et al. 2006; Al-Kaff et al. 2008). Sears (1977) used such crosses between hexaploid wheat cv. Chinese Spring, both in the presence and absence (*ph1b* deletion) of *Ph1*, and the wild relative tetraploid *Aegilops kotschyi* (also termed *Ae. variabilis*), to show that homoeologous COs are induced when the *Ph1* locus is deleted. Interspecific hybrids of the *ph1b* mutant and wild relatives have been subsequently used in plant breeding programmes for introgression purposes (Sears 1977). Sears (1977) observed one rod bivalent at metaphase I in the presence of *Ph1*, and 6.35-7.28 rod bivalents in these *Ph1* absent hybrids. Chiasma frequency in the hexaploid wheat-Ae. *kotschyi* or hexaploid wheat-Ae. *variabilis* hybrids, was between 1-3 in the presence of *Ph1*, and 11-14 in the absence of *Ph1* (Sears 1977; Farooq et al. 1990; Fernández-Calvín and Orellana 1991; Kousaka and Endo 2012).

In this study, both *Tazip4-B2* mutant Cadenza lines, as well as a wild type Cadenza (*TaZIP4-B2*), were crossed with *Ae. variabilis*. The frequency of univalents, bivalents, multivalents and total chiasma frequency was scored at meiotic metaphase I in the resulting F1 hybrid (Fig. 2). In these hybrids, there were similar numbers of rod bivalents to that reported by Sears (1977), with 6.75 (SE 0.17) (Cadenza1691) and 6.64 (SE 0.18) (Cadenza0348) rod bivalents at metaphase I in the *Tazip4-B2* mutants, and 1.48 rod bivalents (SE 0.12) at metaphase I in the wild type Cadenza. Moreover, the chiasma mean frequency was 1.48 (SE 0.12) in the presence of *Ph1* (*TaZIP4-B2*), and 12.21 (SE 0.19) and 12.23 (SE 0.20) in the Cadenza1691-*Ae. variabilis* and Cadenza0348-Ae. *variabilis* hybrids, respectively. The observed chiasma frequencies at metaphase I, in the two *Tazip4-B2* mutant line-*Ae. variabilis* hybrids, are similar to those previously reported at metaphase I in the *Ph1* deletion mutant (*ph1b*)-*Ae. variabilis* hybrids. Thus, the data indicate that *TaZIP4-B2* within the *Ph1* locus is likely to be involved in the suppression of homoeologous COs.

**Fig. 2.**
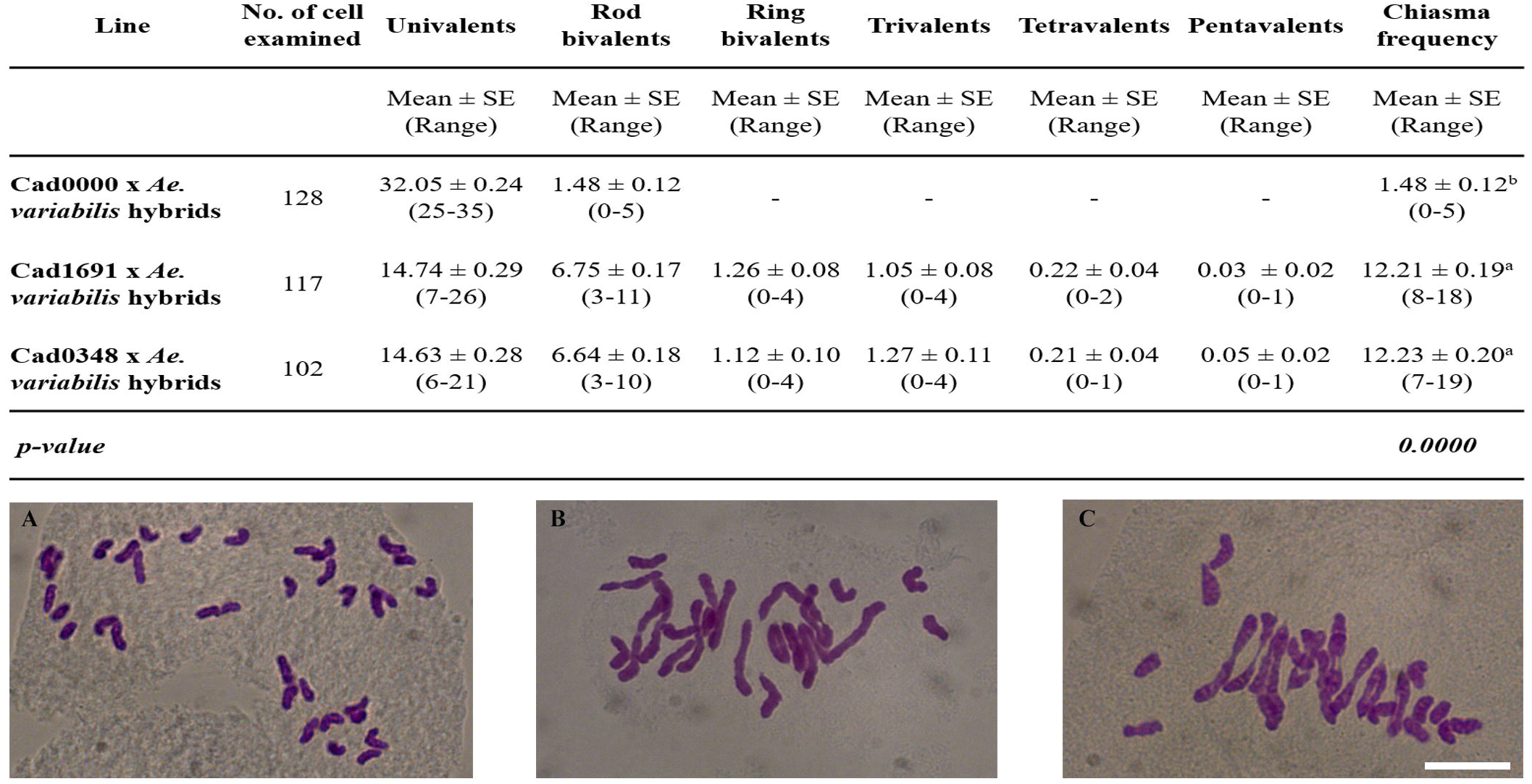
Homoeologous CO frequency at metaphase I is increased in *Tazip4-B2* mutant line-Ae. *variabilis* hybrids (**b** and **c**), in comparison to the wild type Cadenza-Ae. *variabilis* hybrid (**a**). The number of univalents, bivalents, multivalents and chiasma frequency were scored at meiotic metaphase I in Cadenza0000 (*TaZIP4-B2*) x *Ae. variabilis* hybrids, and in Cadenza1691 (*Tazip4-B2*) x *Ae. variabilis* hybrids and Cadenza0348 (*Tazip4-B2*) x *Ae. variabilis* hybrids. The same letter indicates no differences between *TaZIP4-B2* (**a**) and *Tazip4-B2* hybrids (**b** and **c**) in metaphase I at P < 0.05. *Scale bar* represents 10 μm for all panels.

### *Tazip4-B2* mutant Cadenza lines show no multivalents

The frequencies of meiotic associations at metaphase I in hexaploid wheat and the *Ph1* deletion mutant (*ph1b*) have been reported previously (Martín et al. 2014). Martín et al. (2014) observed 20 ring bivalents and one rod bivalent, with a chiasma frequency of 40.97 in the presence of the *Ph1* locus. However, the number of ring bivalents decreased to 14.83, with a reduced chiasma frequency of 35.78, while the number of rod bivalents, univalents, trivalents and tetravalents increased to 4.73, 0.80, 0.20 and 0.37, respectively, when the *Ph1* locus was absent. The number of univalents, bivalents, multivalents and chiasma frequency at meiotic metaphase I was also scored in both the *Tazip4-B2* mutant lines and in the wild type Cadenza (Fig. 3). The *Tazip4-B2* mutant lines exhibited a reduction in the number of ring bivalents at metaphase I, and a slight increase in the number of rod bivalents, from a mean of 1.30 (SE 0.17) in the wild type Cadenza, to 3.29 (SE 0.16) in Cadenza1691 and 3.63 (SE 0.18) in Cadenza0348. This indicates a slight reduction in homologous COs in these *Tazip4-B2* mutant lines. CO frequency was a mean of 40.50 (SE 0.21) in the wild type Cadenza, 38.13 (SE 0.20) in Cadenza1691 and 37.30 (SE 0.23) in Cadenza0348. These observed chiasma frequencies at metaphase I in the two mutant Cadenza lines, are again similar to those previously reported at metaphase I in wheat in the absence of the *Ph1* locus. However, no multivalents were observed, and there was no significant increase in the number of univalents at metaphase I in the *Tazip4-B2* mutant lines, as is normally observed in *Ph1* deletion mutants (Roberts et al. 1999).

**Fig. 3.**
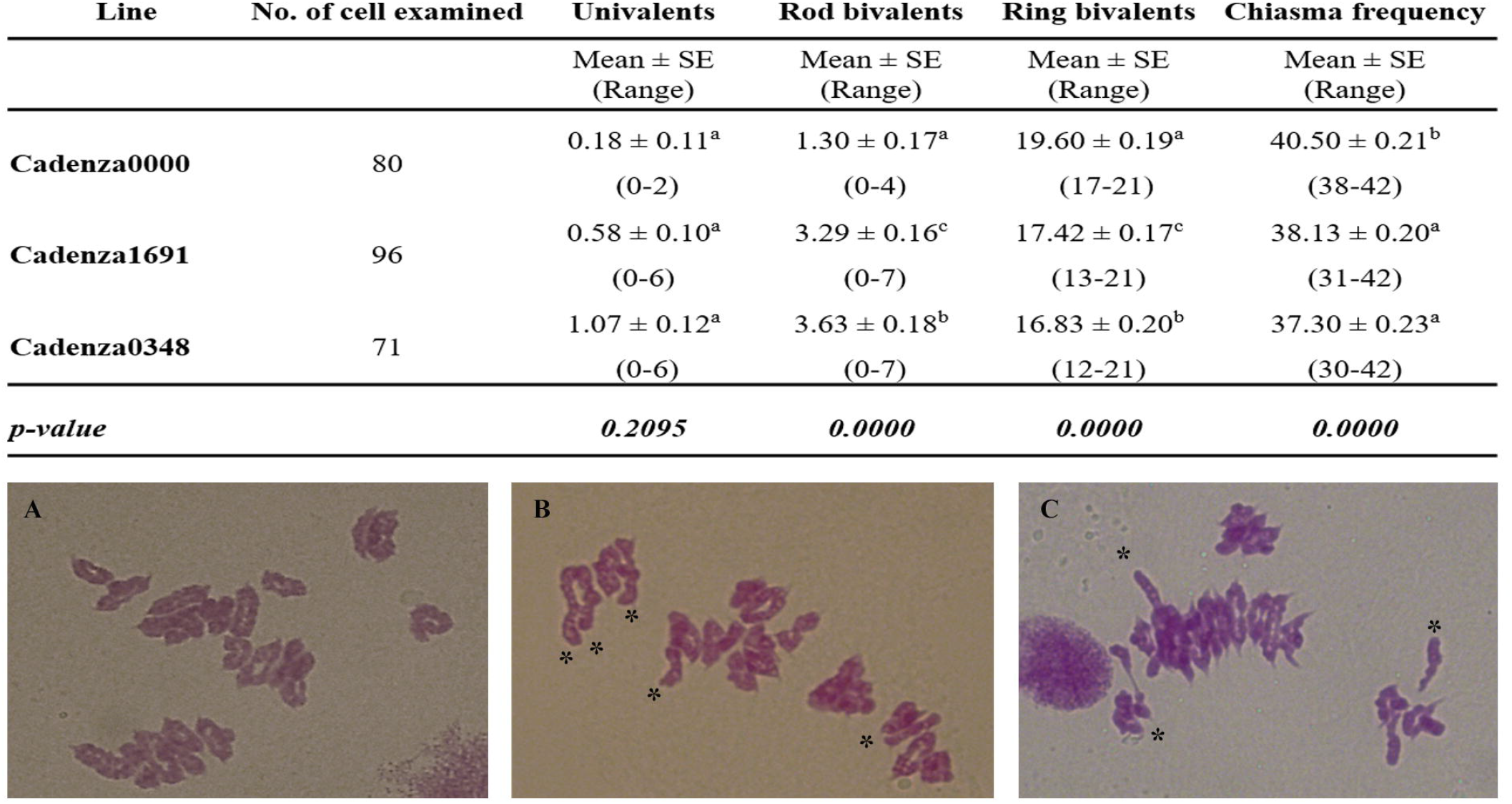
Homologous CO frequency is reduced in *Tazip4-B2* mutant Cadenza lines (**b** and **c**), in comparison to wild type Cadenza (**a**). The number of univalents, bivalents and chiasma frequency were scored at meiotic metaphase I in Cadenza0000 (*TaZIP4-B2*), and in Cadenza1691 (*Tazip4-B2*) and Cadenza0348 (*Tazip4-B2*). Asterisks indicate the presence of rod bivalents in both mutant Cadenza lines. The same letter indicates no differences between *TaZIP4-B2* (**a**) and *Tazip4-B2* wheat (**b** and **c**) in metaphase I at P < 0.05. *Scale bar* represents 10 μm for all panels.

If *Tazip4-B2* mutants had enabled homoeologues to synapse while failing to CO, then a significant increase in univalents would be expected, but this was not observed. This suggests that homoeologous synapsis may not be significantly affected by *TaZIP4-B2*. On the other hand, the lack of multivalents at metaphase I suggests that both mutant lines will exhibit a reduced level of homoeologous exchange or chromosome translocation to that observed in the CS *ph1b* mutant. The *ph1b* mutant line has been reported to accumulate extensive background translocations over multiple generations due to homoeologous synapsis and COs (Sánchez-Morán et al. 2001). Thus, the apparent lack of multivalents in the *Tazip4-B2* mutant lines could allow their exploitation for introgression purposes during plant breeding programmes, rather than the current *ph1b* line.

In summary, seven lines carrying mutations within the *TaZIP4-B2* gene were screened for a phenotype with reduced homologous crossover at metaphase I. Of these, two lines were identified with this phenotype, one carrying a nonsense mutation within *TaZip4-B2*, and the other carrying a mutation in one of the key functional domains of *TaZip4-B2*. When crossed with *Ae. variabilis*, both of these lines also exhibited increased homoeologous crossover at metaphase I in the resulting hybrid, suggesting that the two phenotypes were linked. Therefore, in this case, lines with increased homoeologous crossover had been identified without an initial screen for the desired phenotype. Until now, the only way by which homoeologous crossover at metaphase I could be increased to this extent in wheat-wild relative hybrids was by deletion of the *Ph1* locus, defined as a deletion effect phenotype specific to chromosome 5B. However, an alternative way of reproducing the *Ph1* deletion effect would be to use EMS treatment to generate nonsense or truncation mutations in the homoeologous crossover-suppressing gene within the *Ph1* locus. Analysis of the 1200-line Tilling population revealed that in any given mutant line, 1.5% of genes will have a truncation allele and 2% a missense allele (Krasileva et al. 2017). Thus, the probability of two mutant lines both sharing a truncation or missense mutation by chance in the same second gene is P < 0.0005. The probability that two mutated genes will be located on the same chromosome is extremely low (2.4 x10^−5^), and the probability that they will both be located within the *Ph1* region is even lower (2.4 x10^−7^). Thus, it is extremely unlikely that the increased homoeologous crossover phenotype found in both the two *Tazip4-B2* mutant lines results from a nonsense mutation in a further gene independently linked with *Tazip4-B2* within the *Ph1* locus. These mutants were identified through a non-GM route, and can be exploited as an alternative to the CS *ph1b* mutant. Seeds for both mutants have been deposited with the Germplasm Resource Unit at the John Innes Centre (<u>www.jic.ac.uk/research/germplasm-resources-unit</u>). The accession number of seeds for Cad0348 is W10336 and for Cad1691 is W10337. The seeds for both lines are available on request, free of intellectual property restrictions.

## Author contribution statements

These authors made the following contributions to the manuscript: M-DR, ACM, PS and GM designed the research; M-DR, ACM and JH performed the research; M-DR, ACM, JH, PS and GM analysed the data; M-DR, PS and GM wrote the manuscript.

## Compliance with ethical standards

This research does not involve human participants or animals.

## Acknowledgements

This work was supported by the UK Biotechnology and Biological Research Council (BBSRC), through three grants (Grant BB/J004588/1; Grant BB/M009599/1; Grant BB/J007188/1); and by a Marie Curie Fellowship Grant (H2020-MSCA-IF-2015-703117).

## Supplementary material

**Supplementary Fig. 1** Alignment of coding DNA (**a**) and amino acid sequences (**b**) from *TaZIP4-B2* (TRIAE_CS42_5BL_TGACv1_404600_AA1305800), *TaZIP4-B1* (TRIAE_CS42_3B_TGACv1_225572_AA0809860), *TaZIP4-A1* (TRIAE_CS42_3AL_TGACv1_195180_AA0645950) and *TaZIP4-D1* (TRIAE_CS42_3DL_TGACv1_251716_AA0884100) CS+ refers to wild type and CS- to the *ph1b* mutant.

**Supplementary Fig. 2** Alignment of *TaZIP4-B2* splice variants in wild type Cadenza.

**Supplementary Fig. 3** Alignment of *TaZIP4-B2* coding DNA sequences from wild type Cadenza (WTC) and mutant Cadenza lines (MCL). Both missense and nonsense mutants are highlighted. Primer sequences used to follow mutated genes during crossing are underlined. All primers are shown in direction 5´ → 3´.

**Supplementary Table 1** Detailed information of all transcripts obtained by Kallisto and the statistical analysis of transcripts per million (TPM) in wheat chromosomes 3AL, 3B, 3DL and 5BL, both in presence (WT) and in absence (*ph1b* deletion) of the *Ph1* locus. Data represent mean values ± standard error (SE) from RNA samples collected at late leptotene-early zygotene stage in WT and in *ph1b* deletion.

**Supplementary Table 2** Detailed information on the seven EMS mutant lines selected as possessing potentially interesting mutations within *TaZIP4-B2* (Traes_5BL_9663AB85C.1). The two mutant lines (Cadenza1691 and Cadenza0348) which showed reduced number of COs in Cadenza mutant lines are indicated in bold.

